# CHPF boosts proliferation and invasion of clear cell renal cell carcinoma

**DOI:** 10.1101/2023.06.09.544296

**Authors:** Yaoyu Zhang, Xiaowei Li, Shu Ran, Youguang Zhao, Yuanshun Huang, Tingting Zhou, Qiwu Wang, Jiwen Liu, Qimin Zhang, Xiaodong Li, Taiping Leng, Yihang Luo, Shadan Li

## Abstract

**Background:** Clear cell renal cell carcinoma (ccRCC) is a malignant tumor most commonly seen in the urinary system, featuring quick progression, invasive behavior, frequent recrudesce, and bad prognosis, whereas Chondroitin polymerizing factor (CHPF) is an essential glycosyl transferase of biosynthesis involved chondroitin sulphate. But the relationship between the two has not been fully understood so far. The present research will probe into the relationship between CHPF and ccRCC.

**Methods:** Explore the CHPF expression level in renal ccRCC tissues through bioinformatics analysis of The Cancer Genome Atlas (TCGA) and the real-time quantitative polymerase chain reaction detection system (qPCR); acquire relevant clinical data from the TCGA data-base and use Kaplan-Meier survival analysis to verify the relevance of CHPF expression to the clinical prognosis of ccRCC patients; then, effectively silence CHPF in ccRCC 786-O cells by a lentivirus-mediated approach; next, observe the effects of CHPF on tumor cell proliferation, cell cycle progression, and apoptosis.

**Results:** In ccRCC tissues, CHPF expression has been significantly upregulated; the higher expression, the shorter survival of ccRCC patients. CHPF downregulation in the 786-O cell can effectively hold back the proliferation, apoptosis and cycle of cancer cells.

**Conclusion:** Based on the above results, we arrive at a conclusion that CHPF expression contributes markedly to the development of human ccRCC cells. Therefore, CHPF may act as a potential prognostic marker of ccRCC and provide a new target for ccRCC treatment.

**Simple Summary:** Chondroitin polymerizing factor (CHPF) is an important glycosyltransferase involved in the biosynthesis of chondroitin sulfate, which is thought to have some pro-carcinogenic effects. However, the role and mechanism of CHPF in clear cell renal cell carcinoma (ccRCC) have not been reported. The aim of this study was to investigate the relationship between CHPF and ccRCC. In this study, we found that CHPF was highly expressed in ccRCC and knocking out CHPF can greatly inhibit the proliferation and cell cycle progression of ccRCC and accelerate its apoptosis, suggesting that CHPF can predict the prognosis of ccRCC and a potential target for treatment.

## Introduction

As one of the commonest malignant neoplasms of the urinary system, kidney cancer constitutes 3% of human tumors, with clear cell renal cell carcinoma (ccRCC) being the dominant renal carcinoma sub-type, making up 75% of primary renal cell carcinomas (1,2). Approximately 20% to 30% of ccRCC have metastasized before diagnosis, though the diagnostic results have been improved owing to the progress in imaging technology. In addition, ccRCC is insensitive to chemo- and radio-therapies, so it has an extremely poor prognosis and treatment strategies are limited (3-5). Despite that the mechanisms of tumor formation and development have been extensively studied, the etiology and carcinogenesis of ccRCC are still unknown (6). So, there is a necessity to cut down kidney cancer-associated fatality by a new treatment strategy, like the therapy for specific molecular target.

Chondroitin polymerizing factor (CHPF) is a type II membrane-spanning protein of 775 amino acids, which is vital for chondroitin polymerization activities (7,8). The CHPF gene exists in the 2q35-q36 region of human chromosomes, spans over four exonic areas, and takes a crucial part in cellular functions (9). Above all, CHPF is aberrantly upregulated in multiple types of cancer, and has been proven to be a potential tumour-initiator of ovarian epithelial carcinoma (10), squamous cell carcinoma of the head and neck (SCCHN) (11), and glioma (12), Still and all, what role CHPF plays in ccRCC has so far not been reported and remains almost unknown.

CHPF is highly expressed in human ccRCC tissues and three ccRCC cell lines have been substantiated by our research. We therefore constructed a lentiviral vector (LV-shRNA-CHPF) mediating RNAi targeting of CHPF and further explored in vitro how CHPF facilitates the proliferation and invasion of ccRCC.

## Materials and Methods

### Data Acquisition

The genetic transcription and clinical data of ccRCC were downloaded from The Cancer Genome Atlas (TCGA) data-base (https://portal.gdc.cancer.gov/repository), including a total of 539 tumor samples and 72 normal tissue samples, between which the differences in CHPF expression were analyzed. According to the median expression value, these samples were classified into high and low expression groups, the survival time of which was examined by means of survival analysis or Kaplan-Meier.

### Cancer Cell Lines and Cell Culture

Human renal carcinoma cell lines ACHN, 786-O and Caki-1 were supplied from Shanghai Cell Bank (Shanghai, China), all cultured in DMEM media (Gibco, Carlsbad, CA, USA), each containing 10% fetal bovine serum, in an incubator with 5% CO_2_ at 37°C.

### RNA Isolation and Real-Time Quantitative PCR (qRT-PCR)

Total RNA was extracted with TRIzol -- a reagent purchased from Invitrogen, Shanghai -- from three cell lines according to the instruction book. M-MLV, a reverse transcriptase produced by Promega (USA), was used to synthesize cDNA. The CHPF primers bought from RiboBio Co. Ltd. (a Chinese firm based in Guangzhou) have the following sequences: CHPF upstream primer--5’-GGAACGCACGTACCAGGAG-3’, CHPF downstream primer--5’-CGGGATGGTGCTGGAATACC-3’; GAPDH upstream primer--5’- TGACTTCAACAGCGACACCCA-3’, GAPDH downstream primer--5’- CACCCTGTTGCTGTAGCCAAA-3’. Then, we took 2μg total RNA as a template to perform quantitative PCR (reaction conditions: 45 amplification cycles at 95°C for 5 sec and 60°C for 30 sec) in the Applied Biosystems 7300 fluorescence quantitative PCR system (Applied Biosystems, Foster City, CA, USA) according to the steps in the instruction book of SYBR PrimeScript RT-PCR kit (Takara). We used 2-ΔΔc to indicate the relative gene expression level of each sample.

### Construction of shRNA Lentiviral Vectors (LV) and Cell Transfection

First, we designed a shRNA interference target sequence and cloned it to the pGCSIL-GFP lentivirus with a AgeI/EcoRI site to form a recombinant lentiviral shRNA expression vector through the lentivirus expressing short hairpin RNA (shRNA) targeting the CHPF gene sequence (CTGGCCATGCTACTCTTTG) (Shanghai GeneChem Co.) and the negative control (NC) (TTCTCCGAACGTGTCACGT) (Shanghai GeneChem Co.). Next, we purified lentiviral particles via ultracentrifugation and determined lentivirus titers via end-point dilution. Next, we infected 786-O ccRCC cells with shRNA-CHPF-lentivirus (shCHPF) or NC lentivirus (shCtrl). Then, we inoculated 786-O cells in a 6-well culture plate using a concentration of 40% and 2.0×105 cells/well. 72h after later, we observed the cultured cells with a fluorescence microscopy (MicroPublisher 3.3; Olympus). The last step was collecting the cells to test silencing efficiency by means of quantitative RT-PCR.

### Western Blot

Take cells within 48h of transfection for cytolysis in RIPA buffer (Beyotime, China). Extract proteins, and determine protein concentration via a BCA protein quantification kit (Beyotime, China). Electrophorese the cell lysis solution on SDS-PAGE gel, and transfer to a PVDF membrane bought from Millipore in USA. Notes: The first-used antibody included a polyclonal mouse anti-CHPF (1:1000 dilutions; ABACM, Cambridge, MA, USA). and anti-GAPDH antibody (1:2000 dilutions; Santa Cruz Biotechnology, USA); the antibodies used later were Anti-Rabbit IgG (Cell Signaling Technologies, USA) and Anti-Mouse IgG (Cell Signaling Technologies, USA), which were bound to horseradish peroxidase and diluted at 1:2000. Enhance chemiluminescence using ECL-PLUS/Kit (ThermoFisher Scientific, USA) reagents. Then, study the image results.

### Cell Growth and MTT

After infection with lentivirus, digest the 786-O cells in logarithmic phase with trypsin, then inoculate at 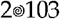 cells/well in 96-well tissue culture plates at 37°C. Starting the day after cell plating, measure the cell quantity continuously using Celigo, an image cytometer imported from Nexcelom Bioscience, Lawrence, MA, USA, count the daily growth rate, and draw the cell survival curve. For the detection of in vitro cell proliferation, use the MTT cell proliferation and cytotoxicity assay kit. Wash the cells with phosphate buffered saline at the end of incubation period according to the instruction book. Then, add MTT, a reagent recommended by the manufacturer. Next, evaluate the absorbance by an enzyme marker at 490 nm to calculate the cellular activity ratio. Repeat each experiment three times.

### Cell Apoptosis and Cell Cycle Assays

Detect cell apoptosis by way of AnnexinV-APC dual-labelling. Collect and dye the cells infected by shCtrl or shCHPF via annexin V-APC as per the manufacturer’s instructions (eBioscience, USA). Measure annexin staining with a flow cytometry namely Calibur II sorter, and analyze cell apoptosis through the software of cell exploration and research (BD Biosciences, USA).

To further confirm changes in the distribution of DNA contents in cells treated with shCHPF, we carried out PI staining: collect the cells group by group and anchor them in 70% ethanol overnight at 4°C. After incubation with 50μg/mL PI and the RNase A (Fermentas International Inc, Canada) in the dark for 30min at 37°C, and carry out FACS analysis as mentioned above. Repeat the above experiments three times.

### Statistical Analysis

Employ SPSS 23.0 to analyze the data and repeat the experiment three times to obtain the final results. Analyze the rawhrough independent samples t-tests. If the difference is P<0.05, it is considered statistically significant.

## Results

### Detection of CHPF mRNA Expression in Renal Cell Carcinoma Lineage Cells

Test the CHPF mRNA expression levels in three renal carcinoma lineage cells (ACHN, 786-O, Caki-1) by RT-PCR. The results show that CHPF is expressed in the three cells (Fig 1 and S1 Table.).

**Fig 1.**
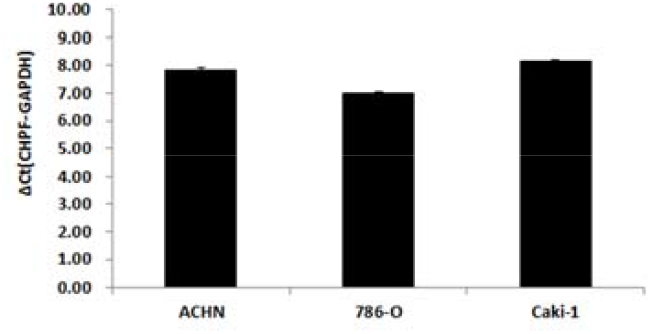
The expression of CHPF in three renal cancer cell lines was detected by quantitative RT-PCR.

### High CHPF Expression in ccRCC Tissues

To verify whether CHPF participates in the development of ccRCC, we made differential analysis using the data from TCGA. The results indicate that CHPF expression increases notably in ccRCC tissues (n=539) than in normal tissues (n=72) (Fig 2A). Then, we divided CHPF expression into high and low groups according to the median of the CHPF expression quantity. The overall survival (OS) of the patients in the TCGA database was analyzed by Kaplan-Meier. The results show that CHPF expression levels have an obvious impact on patient OS, and the OS of ccRCC patients has been remarkably lowered in the high expression group (Fig 2B).

**Fig 2.**
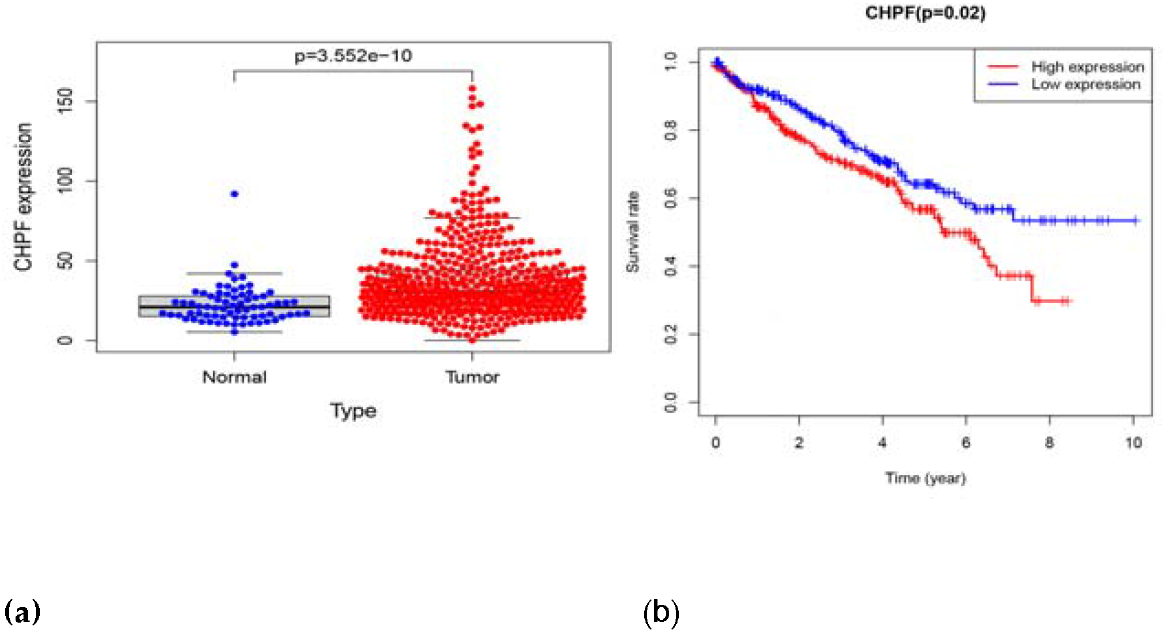
Significance of CHPF expression in the TCGA database. A: CHPF expression in ccRCC tissue and adjacent normal tissue in TCGA. B: CHPF expression and overall survival in ccRCC patients in TCGA cohort.

### Determination of Gene Knockdown Efficiency through Western Blot

Contaminate human ccRCC 786-O cells using shCHPF or shCtrl. Detect the CHPF protein level in the 786-O cell by Western Blot. It is evident from the Western Blot results that in 786-O cell the target has a significant knockdown effect on the endogenous expression of CHPF genes at the protein level (Fig 3).

**Fig 3.**
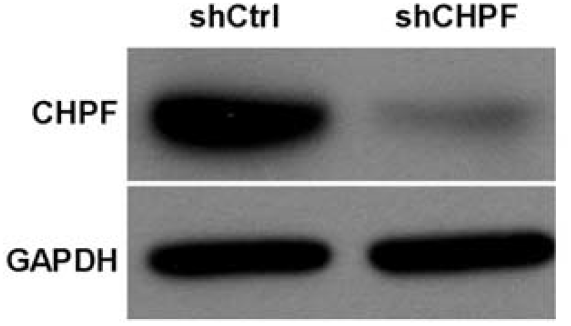
Western blotting-verified transfection efficiency of 786-O cell lines. 3.4. Knockdown of CHPF in Human ccRCC 786-O Cells by shRNA System

To discuss the function of CHPF in ccRCC, we generated shCHPF and shCtrl that expressed Green fluorescent protein (GFP) and infected human ccRCC 786-O cells. The concentration dose of shCHPF was 4×108 TU/mL. In the shCHPF group and the shCtrl group, fluorescence observations reveal that the cells were infected with an efficiency of more than 80% and the cell state was normal (Fig 4A). Its knockdown effect was analyzed by qRT-PCR. After five days of infection, the CHPF mRNA levels in 786-O cells infected with shCHPF (1.001 ± 0.006) were obviously lower than those infected with shCtrl (1.001 ± 0.067) (Fig 4B and S2 Table.).

**Fig 4.**
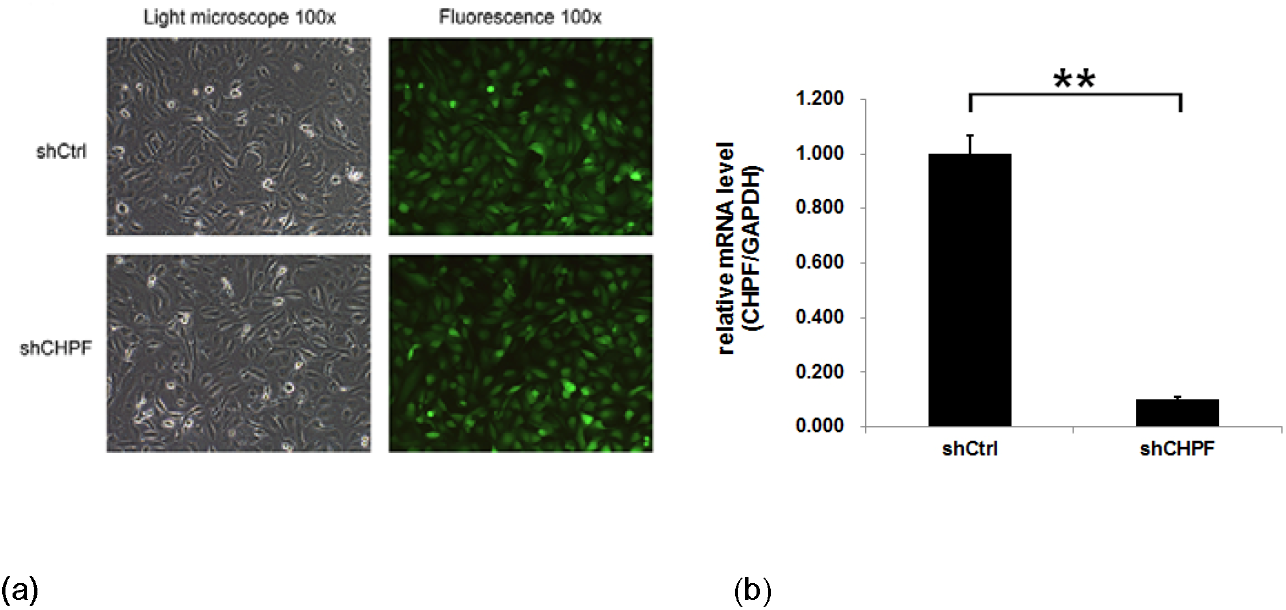
A lentiviral vector expressing shCHPF was constructed and transfected into the glioma 786-O cells. (A) The infection efficiency was examined by fluorescence and light microscopy at 72 h post-infection. Most cells exhibited positive green fluorescence in both the shCHPF and shCtrl groups. (B) The knockdown efficiency of CHPF was assessed by quantitative RT-PCR assay; **P<0.01.

### Knockdown of CHPF to Significantly Restrain the Growth of 786-O Cells

To test how CHPF influences cell growth, we inoculated the 786-O cells infected with shCHPF or shCtrl into 96-well plates three days later for daily cytomic analysis for five straight days. Fig 5 shows the cell quantity in the shCHPF group declines continuously since infection whereas the space between shCHPF and shCtrl groups widens with time (S3 Table.). The research findings indicate CHPF downregulation has remarkably inhibited the growth of 786-O cells.

**Fig 5.**
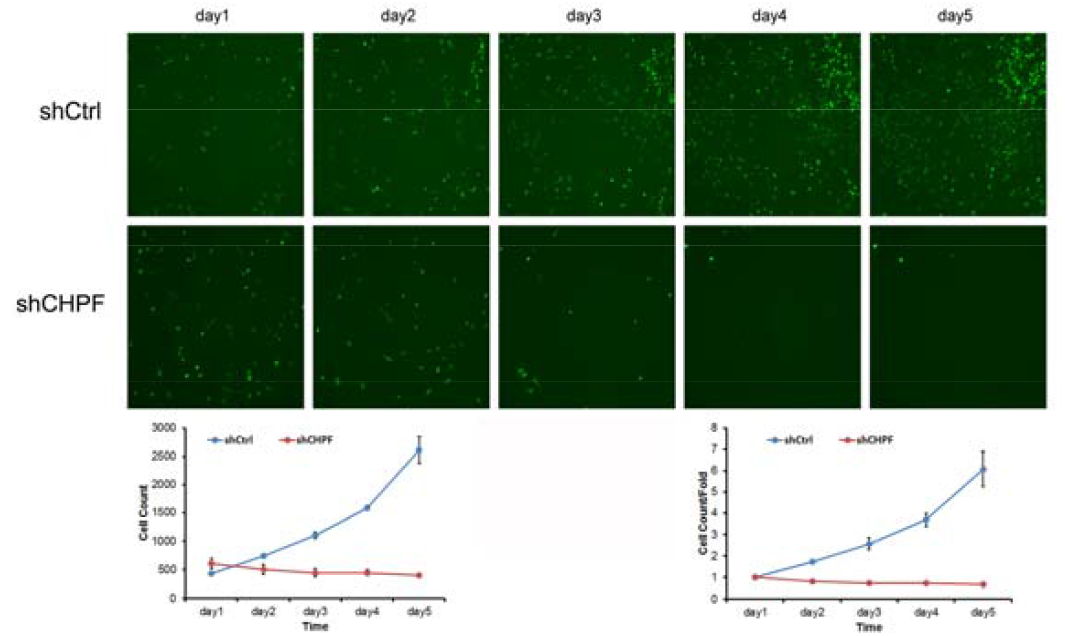
Knockdown of CHPF decreases the proliferation of glioma 786-O cells. High-content cell imaging was applied as indicated daily to acquire raw images of cell growth after lentiviral infection.

To further evaluate the regulation effect of CHPF on ccRCC proliferation, we conducted the MTT assay again. Fig 6 shows the downregulation of CHPF expression has largely reduced the proliferative potential of ccRCC cells (P<0.01). The growth of 786-O cells was largely suppressed after shCHPF treatment compared with the shCtrl group. The in vitro growth of shCHPF cells obviously slowed down from day 4 (shCtrl, 3.606±0.098 vs. shCHPF 1.832±0.044; P<0.001) and day 5 (shCtrl, 4.041±0.073 vs. shCHPF, 1.859±0.094; P<0.001) (S4 Table.). Based on the above, we drew the conclusion that the in vitro growth of 786-O cells is dependent on high expression of CHPF.

**Fig 6.**
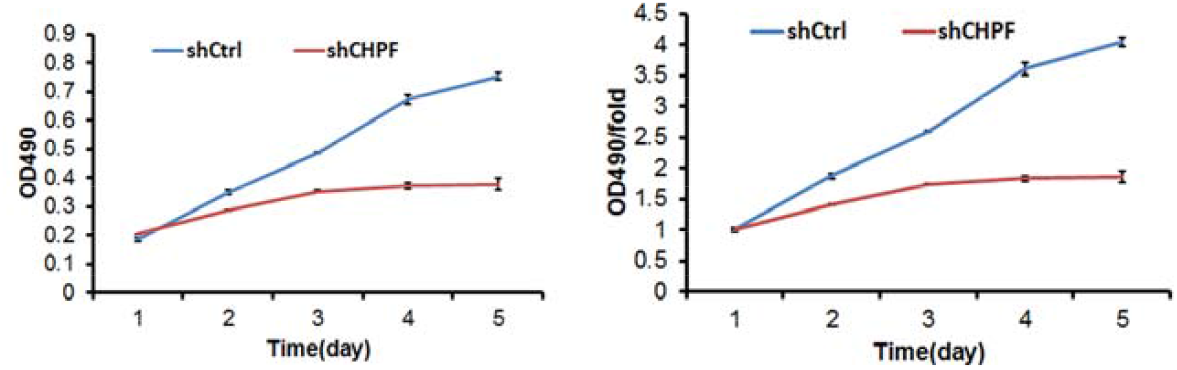
MTT assay displaying the proliferation ability of the 786-O1 cells after transfection with shCHPF.

### CHPF Inhibition to Induce G2/M Phase Arrest in ccRCC 786-O Cells

To verify the impact of CHPF on the cell growth cycle, we detected the 786-O cell cycle by flow cytometry. In Fig 7, the cell cycle in the shCHPF group comprises the following phases: G0/G1, 42.6±0.77%; S, 27±0.34%; G2/M, 30.4±0.44%. The composition of the shRNA-Ctrl group is distributed as G0/G1 phase, 44.22±0.86%; S phase, 35.59±1.01%; G2/M phase, 20.19±0.88%. When CHPF was absent after knockdown, the quantity of cells entering phase G2/M rises up to 29.4% (P<0.001) whereas the quantity of cells in phase S fell 24.14% (P<0.001) (S5 Table.). Taken together, these data indicate that CHPF regulates 786-O cell growth, and particularly blocks cell cycle progression in the G2/M phase.

**Fig 7.**
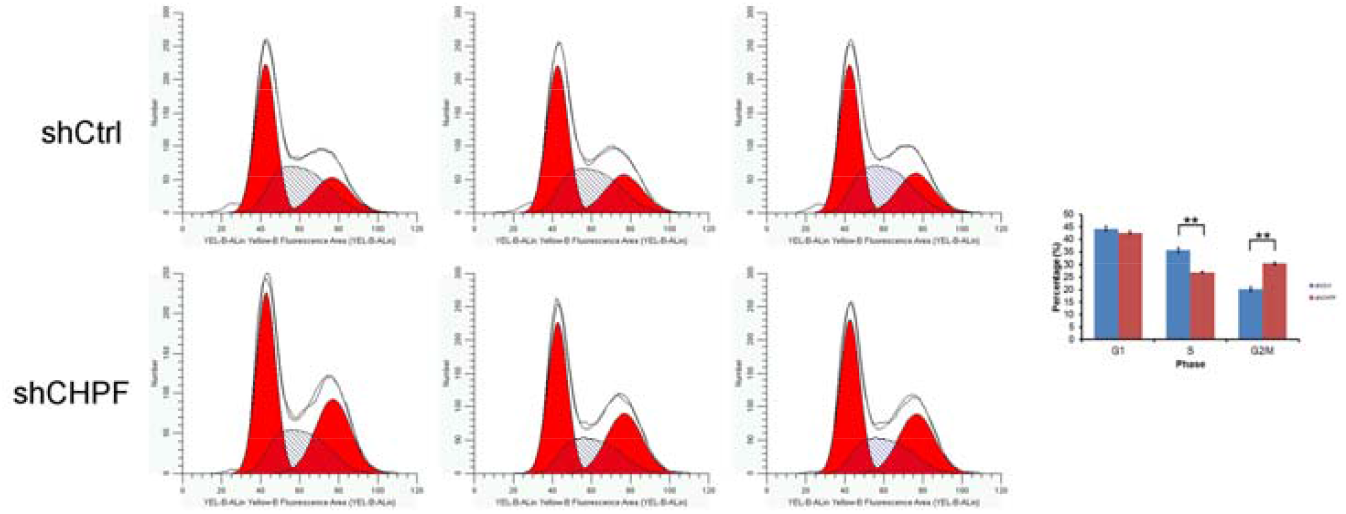
Cell cycle of 786-O cell lines was examined by flow cytometry. The 786-O cells were arrested in G2/M phase after knockdown of CHPF. The experiments were performed in triplicate. The data were shown as mean ± SD. *P < 0.05, **P < 0.01, ***P < 0.001.

### Increase of ccRCC 786-O Cell Apoptosis via CHPF Knockout

We detected the effect of CHPF knockdown on the apoptosis of 786-O cells by flow cytometry. As shown in Fig 8, the cell apoptosis (6.48%±0.26%) of the shCHPF group is higher than that (3.68%±0.11%) of the shCtrl group (P<0.001) (S6 Table.). This indicates remarkable association between CHPF and the apoptosis of 786-O cells. In addition, the cell cycle assay results demonstrate CHPF silencing has stifled 786-O proliferation by inducing cell cycle arrest and apoptosis in the G2/M phase.

**Fig 8.**
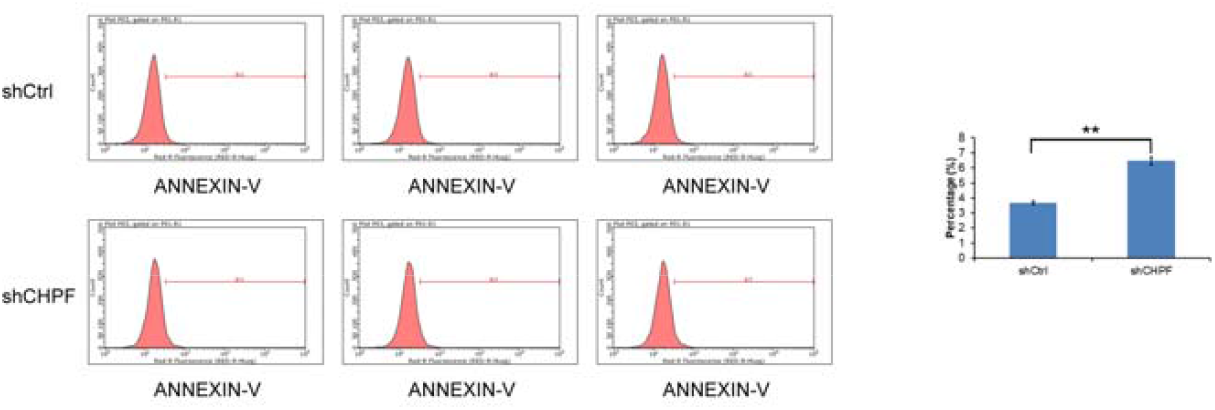
CHPF knockdown promotes glioma 786-O cell apoptosis. The apoptotic rate of the 786-O cells after transfection with shCHPF analyzed using flow cytometry. The experiments were performed in triplicate. The data were shown as mean ± SD. *P < 0.05, **P < 0.01, ***P < 0.001.

## Discussion

ccRCC is a malignancy stemming from the epithelial system of renal parenchyma-forming uriniferous tubules, and its pathogenesis is the result of combined actions of multiple genes (13,14). Moreover, it is difficult to be diagnosed at an early stage and has an undesirable prognosis (15). Therefore, to study the mechanisms of ccRCC occurrence and development is indispensable. Chondroitin sulfate (CS), a polysaccharose comprising repeated disaccharide units of N-acetyl-d-d-galactosamine and d-glucuronide residues, is modified via sulfated residues at different sites. CS biosynthesis and sulfated balance are in strict control and exert a pivotal role in progressive disease (16,17). Previous researches using inhibitors or enzymes to degrade CS chains indicate that CS has a big impact on the metastasis, proliferation and adhesion of tumors (18,19). CHPF is the fifth important member of the chondroitin sulphate polymerase family found to interact with other family members to regulate the extension of the chondroitin sulphate chain (20,21). So, CHPF may facilitate tumor progression. Past researches demonstrate that CHPF is abnormally upregulated in some types of cancer, e.g., colorectal carcinoma (22), laryngocarcinoma (23), and glioma (12), yet the function of CHPF in ccRCC remains unclear.

In our research, we confirmed high expression of CHPF in ccRCC tissues by differential analysis of TCGA database. Kaplan-Meier survival analysis indicate the high expression group has a shorter OS compared to the low expression group. RT-qPCR suggests high expression of CHPF in three renal cancer lines. Subsequently, we constructed a CHPF shRNA expressed LV and transfected it into 786-O cells, which stably down-regulated the expression level of CHPF genes in 786-O cells in vitro. Experiments reveal that CHPF downregulation can inhibit ccRCC progression by slowing down cell growth, increasing the percentage of cell apoptosis, blocking the cell cycle and suppressing cell progression at the G2/M phase. Thus, our research results confirm that CHPF has promoted the growth of ccRCC 786-O cells and the CHPF gene can be used as a target for the treatment of ccRCC. Our research has some limitations too, mainly including insufficient data on follow-up visits with ccRCC patients and unclear downstream regulation mechanism.

## Conclusions

All in all, our research finds that CHPF expression has been upregulated in ccRCC tissues and that high CHPF expression is positively correlated to bad prognosis. Knocking out CHPF can greatly inhibit the proliferation and cell cycle progression of ccRCC and accelerate its apoptosis. According to our research findings, CHPF may contribute to ccRCC formation and development as a tumor-initiator, which can thus be taken as a prognostic indicator or therapeutic target of ccRCC.

## Supporting information

**S1 Table.** The table of RT-PCR data for three renal carcinoma lineage cells.

| Name of cell | GAPDH | CHPF | $\Delta Ct$ | $\Delta Ct$ 均值 | STDEV. |
| --- | --- | --- | --- | --- | --- |
| ACHN | 12.01 | 19.71 | 7.7 | 7.81 | 0.103 |
|  | 11.89 | 19.79 | 7.9 |  |  |
|  | 11.96 | 19.8 | 7.84 |  |  |
| 786-O | 12.62 | 19.68 | 7.06 | 7.01 | 0.046 |
|  | 12.48 | 19.48 | 7 |  |  |
|  | 12.52 | 19.49 | 6.97 |  |  |
| Caki-1 | 12.59 | 20.75 | 8.16 | 8.14 | 0.082 |
|  | 12.47 | 20.68 | 8.21 |  |  |
|  | 12.52 | 20.57 | 8.05 |  |  |

**S2 Table.**
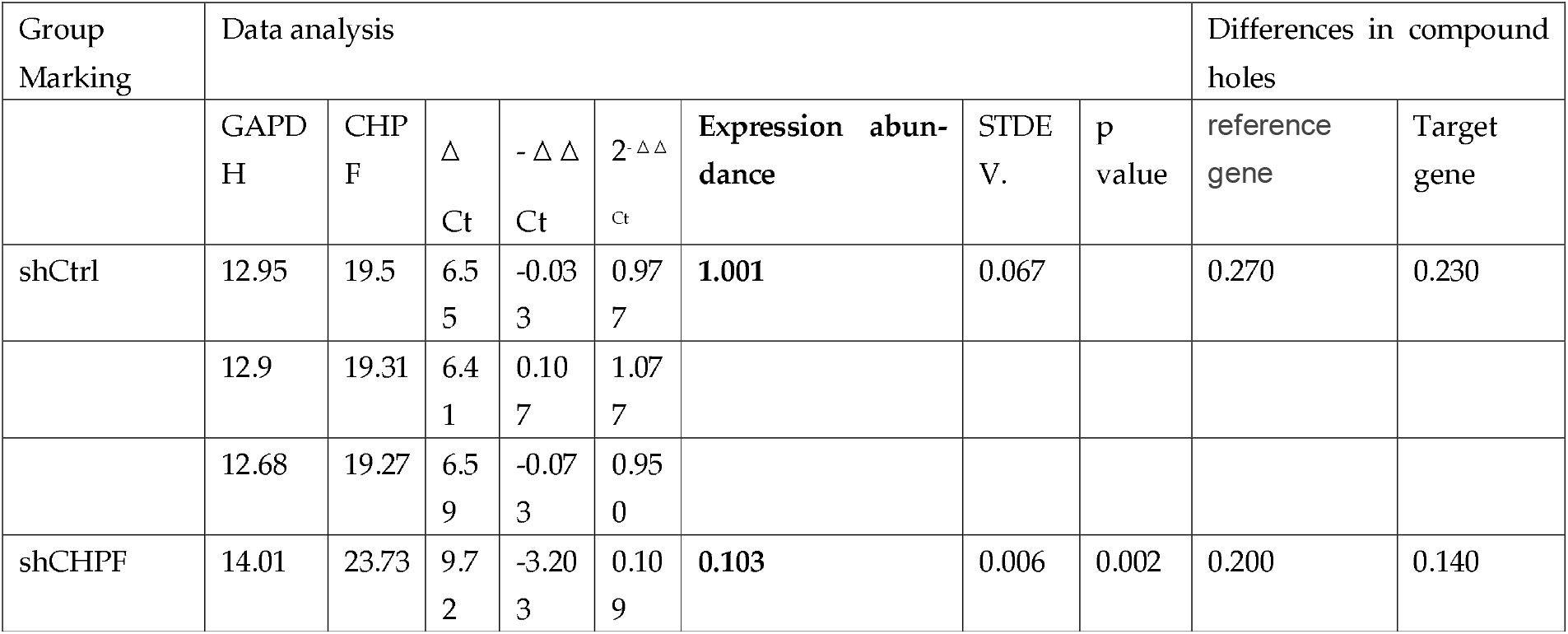
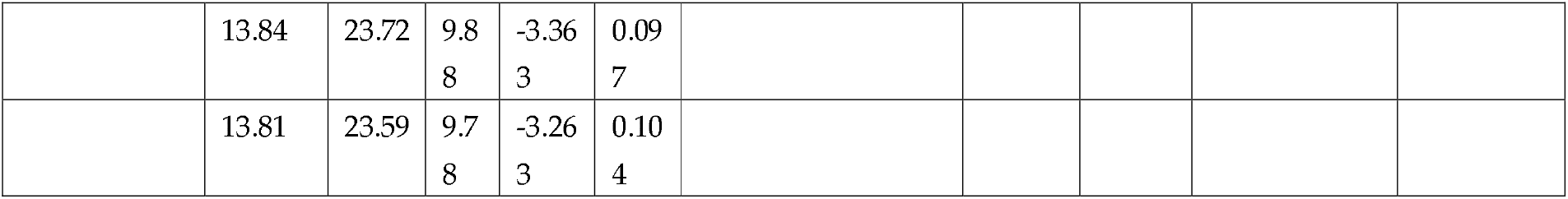
The table of data on target gene knockdown efficiency at the mRNA level by qPCR.

**S3 Table.**
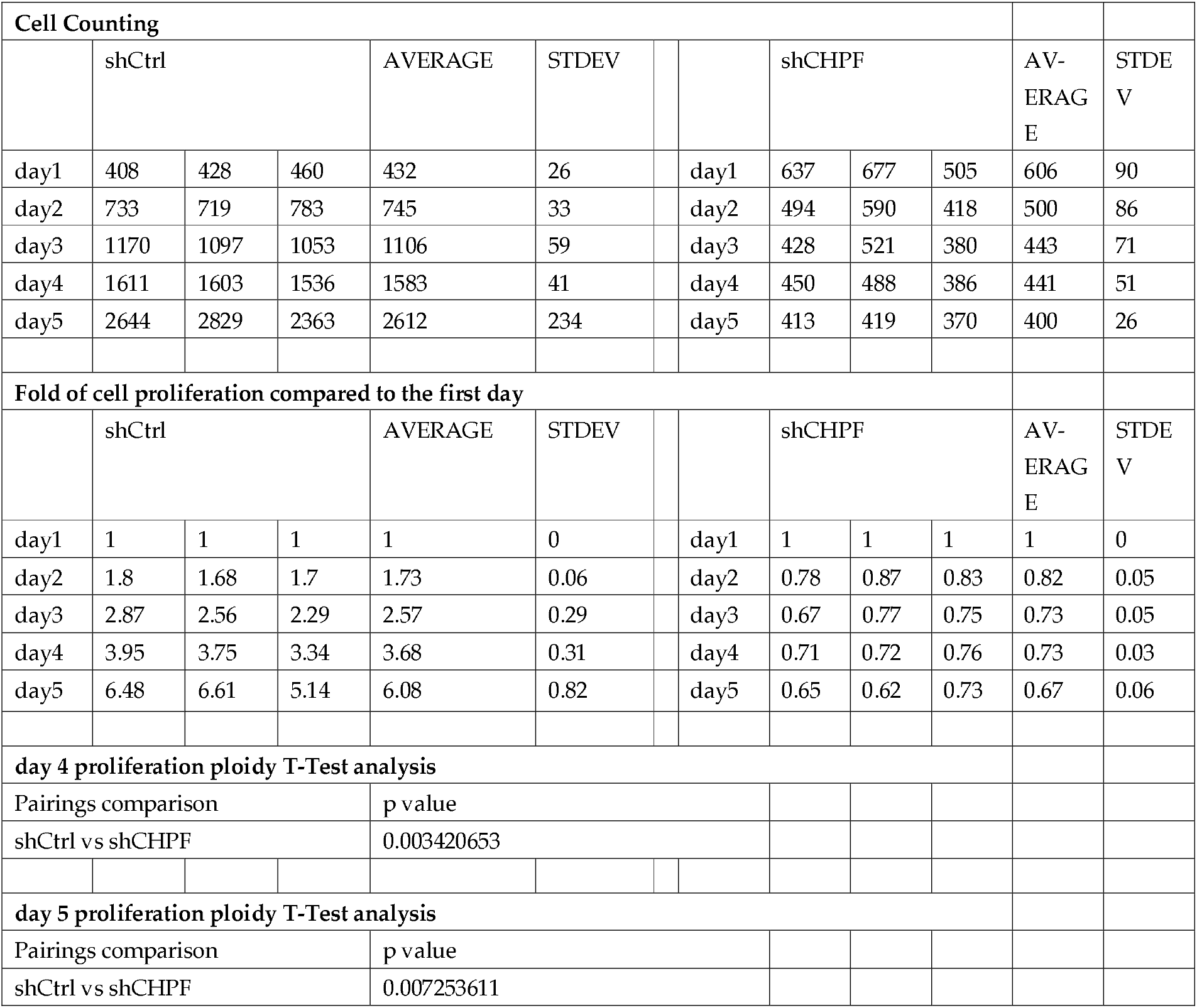
The table of Celigo cell count data for cell growth assays.

**S4 Table.**
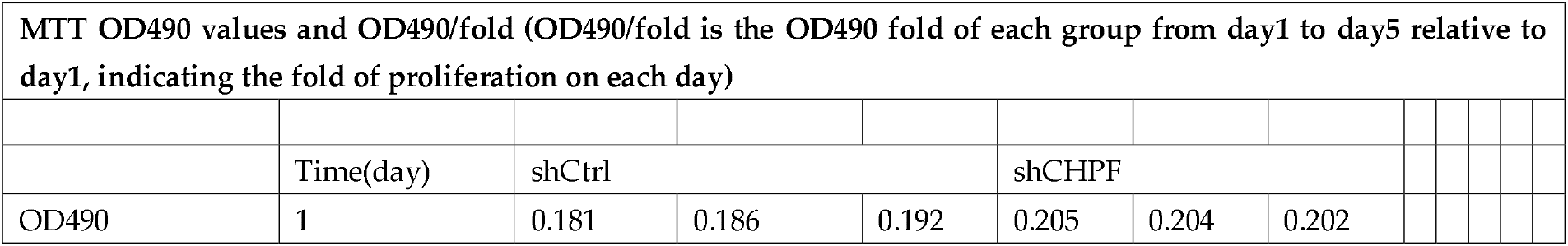
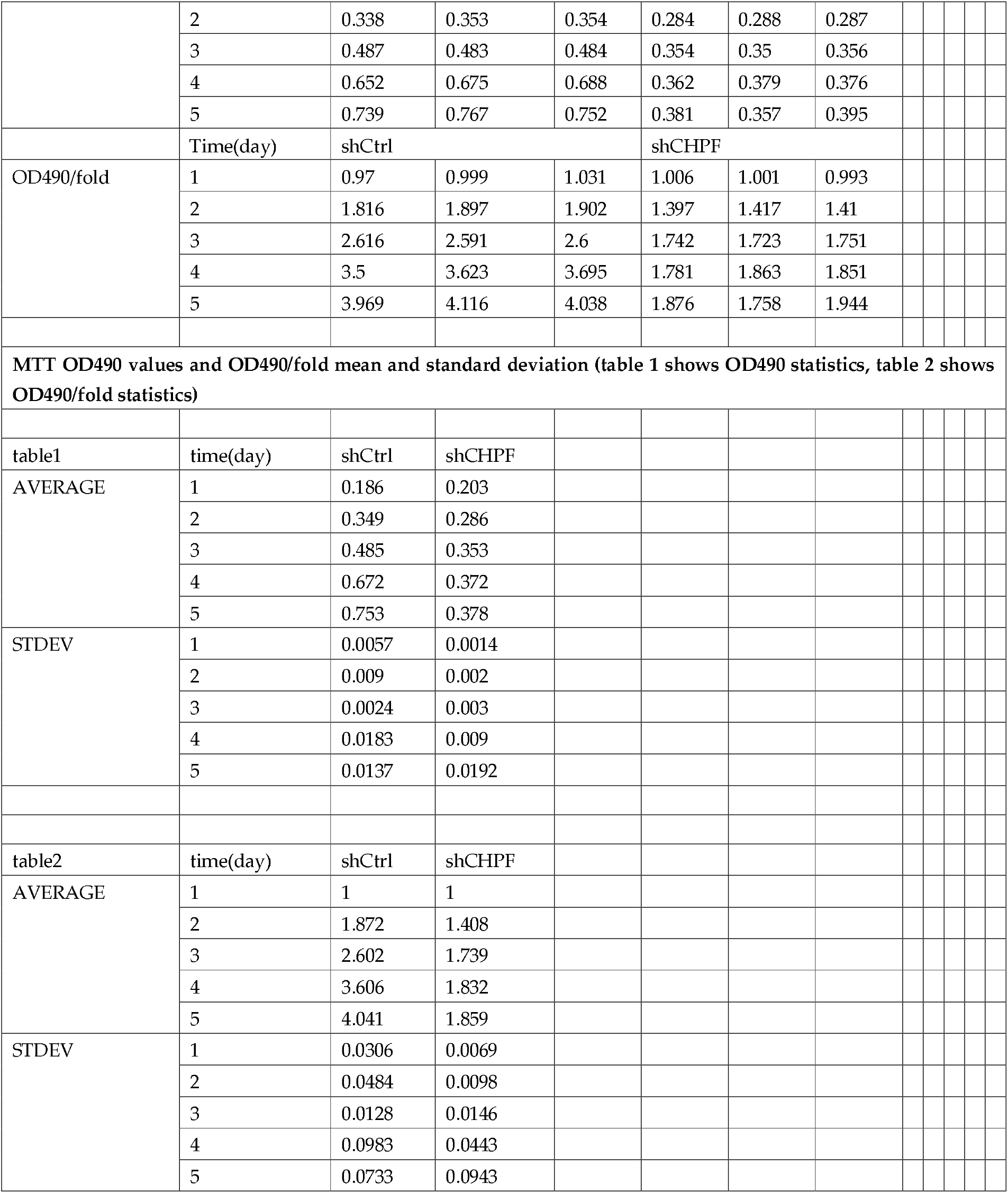
The table of data from the MTT test.

**S5 Table.**
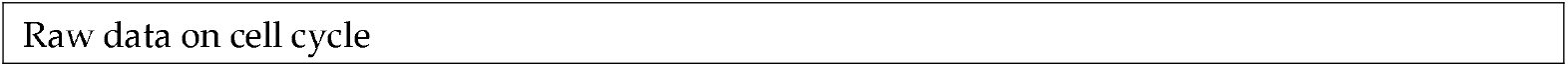
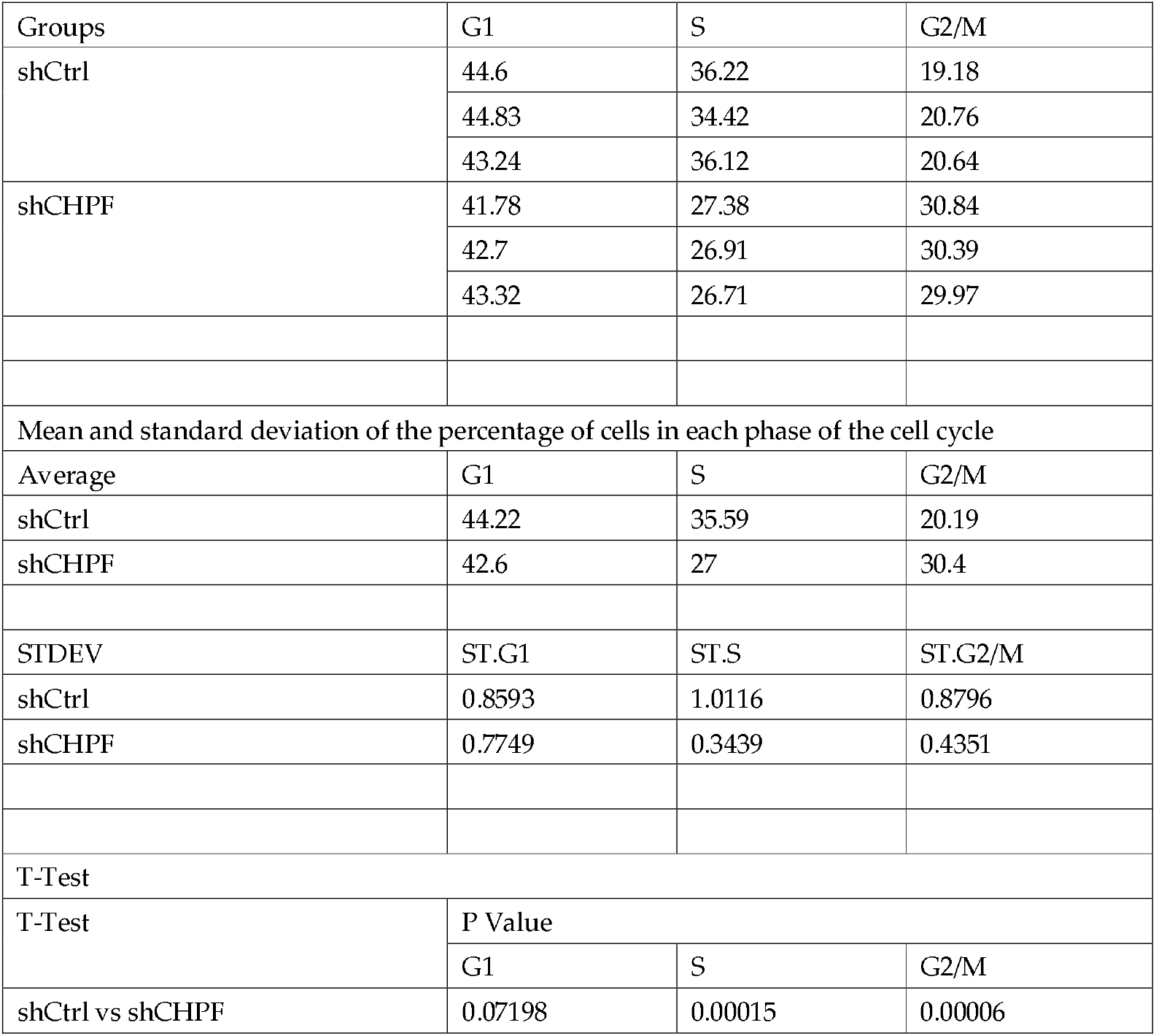
The table of experimental data on the cell cycle.

Raw data on cell cycle
| Groups | G1 | S | G2/M |
| --- | --- | --- | --- |
| shCtrl | 44.6 | 36.22 | 19.18 |
|  | 44.83 | 34.42 | 20.76 |
|  | 43.24 | 36.12 | 20.64 |
| shCHPF | 41.78 | 27.38 | 30.84 |
|  | 42.7 | 26.91 | 30.39 |
|  | 43.32 | 26.71 | 29.97 |
| Mean and standard deviation of the percentage of cells in each phase of the cell cycle |  |  |  |
| Average | G1 | S | G2/M |
| shCtrl | 44.22 | 35.59 | 20.19 |
| shCHPF | 42.6 | 27 | 30.4 |
| STDEV | ST.G1 | ST.S | ST.G2/M |
| shCtrl | 0.8593 | 1.0116 | 0.8796 |
| shCHPF | 0.7749 | 0.3439 | 0.4351 |
| T-Test |  |  |  |
| T-Test | P Value |  |  |
|  | G1 | S | G2/M |
| shCtrl vs shCHPF | 0.07198 | 0.00015 | 0.00006 |

**S6 Table.**
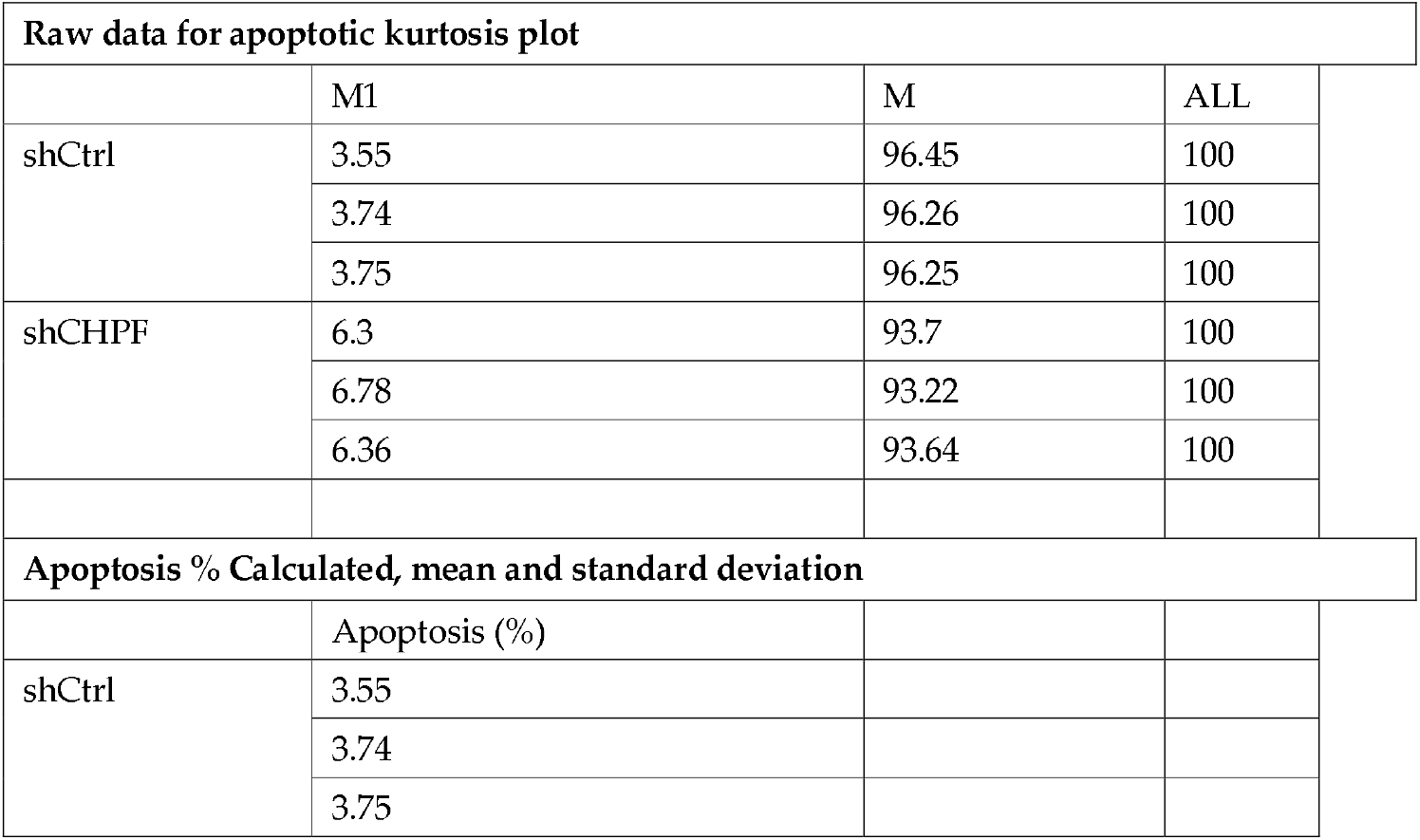
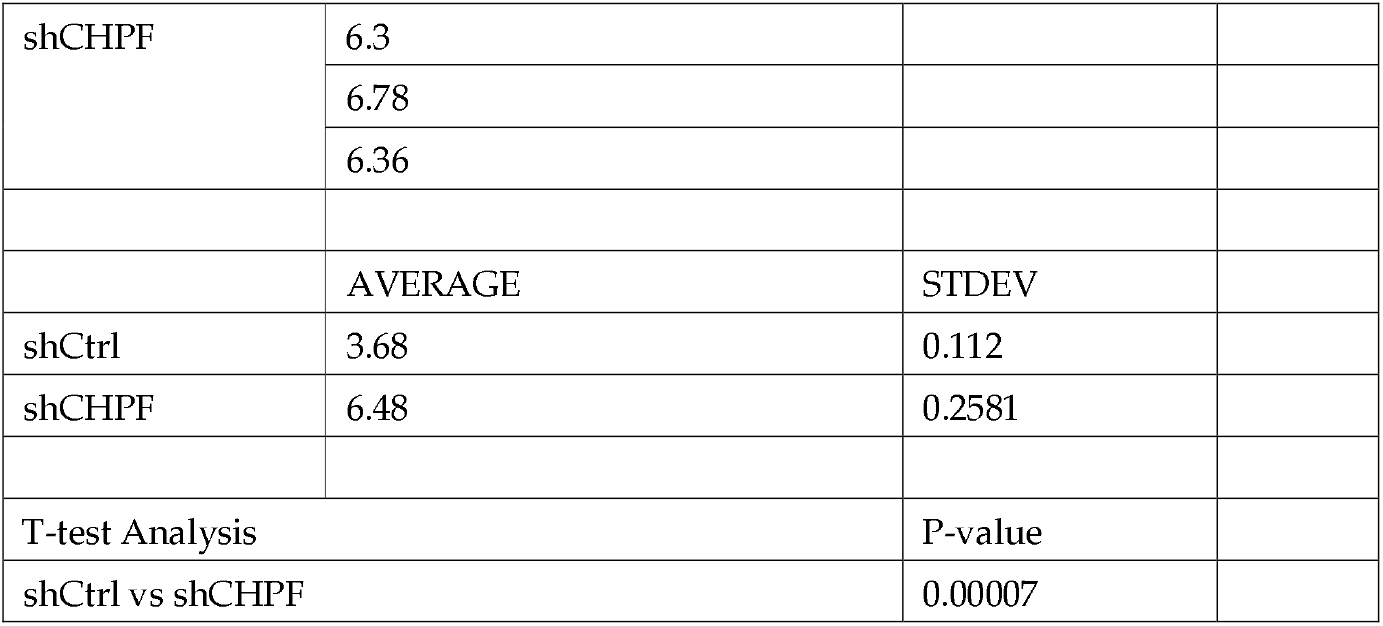
The table of apoptosis assay data.

| Raw data for apoptotic kurtosis plot |  |  |  |
| --- | --- | --- | --- |
|  | M1 | M | ALL |
| shCtrl | 3.55 | 96.45 | 100 |
|  | 3.74 | 96.26 | 100 |
|  | 3.75 | 96.25 | 100 |
| shCHPF | 6.3 | 93.7 | 100 |
|  | 6.78 | 93.22 | 100 |
|  | 6.36 | 93.64 | 100 |
| Apoptosis % Calculated, mean and standard deviation |  |  |  |
|  | Apoptosis (%) |  |  |
| shCtrl | 3.55 |  |  |
|  | 3.74 |  |  |
|  | 3.75 |  |  |

|  |  |  |
| --- | --- | --- |
| shCHPF | 6.3 |  |
|  | 6.78 |  |
|  | 6.36 |  |
|  | AVERAGE | STDEV |
| shCtrl | 3.68 | 0.112 |
| shCHPF | 6.48 | 0.2581 |
| T-test Analysis |  | P-value |
| shCtrl vs shCHPF |  | 0.00007 |

## Author Contributions

SDL, YGZ and YYZ made substantial contribution to the design of this study. The experiments were performed by YYZ, XWL, TTZ and YHL. YYZ performed bioinformatic and statistical analysis. Data analysis was carried out by QMZ, XDL and TPL. YYZ and SDL supervised research and wrote the manuscript. Moreover, WJM and YGZ checked and improved the manuscript and all authors have approved the submission of this manuscript.

## Data Availability Statement

Publicly available datasets were analyzed in this study. This data can be found here: http://portal.gdc.cancer.gov/repository.

## Acknowledgments

We thank Dr. Wenjun Meng from WCHSCU for his language revision during manuscript preparation.

